# Foraging niche partitioning of three *Myotis* bat species and marine fish consumption by *Myotis pilosus* in a subtropical East Asian region

**DOI:** 10.1101/2024.08.26.609698

**Authors:** Xiaodong Wei, Emily Shui Kei Poon, John Chun Ting Chung, David Tsz Chung Chan, Chung Tong Shek, Wing Chi Tsui, Huabin Zhao, Simon Yung Wa Sin

## Abstract

Most bats are insectivorous, but some species have evolved the ability to prey on fish. Although piscivory has been confirmed in the Rickett’s big-footed myotis (*Myotis pilosus*), the extent of piscivory of other cohabiting *Myotis* species is uncertain. This study aims to explore the dietary niches and fish consumption of three *Myotis* species in a subtropical East Asian region, and specifically the fish diet of *M. pilosus*. Our findings reveal, for the first time, that *M. pilosus* consumes marine fishes, in contrast to previous research conducted in inland regions that suggested year-round consumption of cyprinids in freshwater habitats. We also observed seasonal variation in the diets of *M. pilosus*. It predominately hunted wide-banded hardyhead silverside, sailfin flying fish, and shorthead anchovy during the wet season, while mainly preying upon mullets during the dry months. In more inland areas, *M. pilosus* was found to primarily feed on invasive freshwater poeciliids. Furthermore, *M. pilosus* consumed more fish during the dry season, while there was a greater consumption of insects during the wet months. Most notably among our findings is the consumption of fish by two individuals of Horsfield’s myotis (*M. horsfieldii*), indicating that the species is potentially piscivorous. We revealed that both *M. horsfieldii* and *M. pilosus* consumed water striders, suggesting that foraging of aquatic insects could be driving the evolution of fishing behavior. Our findings have also shed light on the flexibility of foraging behavior in piscivorous bats.

## 1. Introduction

Bats are the only mammals capable of sustained flight. They use echolocation during flight to navigate and forage for food nocturnally. There are over 1400 known bat species worldwide, forming a diverse group that have adapted to a wide range of diets (Taylor, 2019). The majority of bats are insectivorous, feeding primarily on insects, while others are frugivorous and nectarivorous bats, which consume fruit and nectar, respectively (Ramírez-Fráncel et al., 2022). In addition to insects, certain bat species have adapted to feed on other animals; for instance, sanguivorous vampire bats consume blood (Riskin and Carter, 2023), and piscivorous bats prey on fish (Aizpurua and Alberdi, 2018). Piscivorous bats can be found among two distinct genera: *Noctilio* (bulldog bats) and *Myotis* (mouse-eared bats). Among these, only the *N. leporinus* (greater bulldog bat) in Latin America has been confirmed as mainly piscivorous (Brooke, 1994). Three other species display ‘limited’ fishing behaviors, including *M. vivesi* (fish-eating myotis) in the Gulf of California (Otálora-Ardila et al., 2013), *M. pilosus* (Rickett’s big-footed myotis) in southern and eastern China, Vietnam, and Laos (Ma et al., 2003), and *M. capaccinii* (long-fingered myotis) in the Mediterranean Sea (Aihartza et al., 2003). Additionally, several species have ‘unconfirmed’ fishing behaviors, such as *N. albiventris* (lesser bulldog bat), *M. daubentonii* (Daubenton’s myotis), and *M. macropus* (large-footed myotis).

The evolution of fishing behavior in bats has long fascinated scientists, and comprehending the dietary compositions of various piscivorous bats is crucial for understanding their foraging ecology and unravelling the intricacies of these evolutionary processes. While fishing behavior is an important hunting strategy, it is worth noting that none of these species are obligate piscivorous (Aizpurua et al., 2014). Fish-eating bats also prey upon arthropods to varying extents, and the prevalence and intensity of fishing or hunting arthropods can be influenced by different factors such as their morphological traits, prey availability, and interspecific competition within their habitats (Chang et al., 2019).

In Asia, *M. pilosus* is relatively large for its genus, exhibiting a body length ranging from 51 to 65 mm (Wilson and Mittermeier 2019). They possess remarkably enormous feet with enlarged, laterally compressed claws, resembling those of *N. leporinus* (Fish et al., 1991, Ospina-Garcés et al., 2016). However, it was not until 2003 that the piscivorous nature of *M. pilosus* was confirmed (Ma et al., 2003). The bats capture insects in mid-air and use their hind limbs to employ a trawling method for catching fish (Jiang et al., 2003). While fish remains have been discovered in their diets in some regions, regular fish consumption has only been verified in Beijing in northern China (Ma et al., 2006). Studies conducted in Beijing, Shandong, and Guizhou have shed light on the dietary habits of *M. pilosus* (Ma et al., 2006, Chang et al., 2019, Ma et al., 2003, Wang et al., 2024). These studies reported on the piscivorous and insectivorous feeding behaviors of *M. pilosus*, with species from the order Cypriniformes (cyprinids) being their primary fish prey. In Beijing, they found that *M. pilosus* consumed grass carp (*Ctenopharyngodon idella*), Eurasian carp (*Cyprinus carpio*), pale chub (*Zacco* spp.), and Amur minnow (*Rhynchocypris* spp.) (Ma et al., 2006, Chang et al., 2019). In Shandong, in addition to grass carp and Eurasian carp, they also found bitterlings (*Rhodeus* spp.) to be their primary prey (Chang et al., 2019). These fish species are similarly found in freshwater habitats within inland areas (Froese and Pauly, 2024). Additionally, *M. pilosus* was found to consume at least seven orders of insects, such as Coleoptera (beetles), Lepidoptera (moths and butterflies), Hemiptera, Diptera, and more (Ma et al., 2006, Chang et al., 2019).

However, discrepancies in the extent of fishing incidence have been observed among these studies. One possible factor contributing to these discrepancies could be the variations in spatiotemporal prey availability across different sampling locations and seasons. Further research on their fish and insect consumption is crucial to gain a better understanding of how resource variation or resource partitioning is associated with its adaptations to the environment through foraging strategy. Thus, it is necessary to investigate the dietary compositions of *M. pilosus* in various locations or habitats across seasons to provide further insights into their fishing behavior. However, beyond the few populations studied, the dietary composition of *M. pilosus* remains largely unknown.

There have been speculations that several other Asiatic *Myotis* species, such as *M. macrotarsus* (pallid large-footed myotis), *M. horsfieldii* (Horsfield’s myotis), *M. hasseltii* (lesser large-footed myotis), *M. stalkeri* (Kei myotis), *M.* adversus (large-footed bat), *M. macrodactylus* (big-footed myotis), and *M. macropus*, might be piscivorous due to their large hind feet and claws (Aizpurua and Alberdi, 2018). Notably, *M. horsfieldii* is a native bat species in South Asia. Although it is smaller compared to *M. pilosus*, with a body length ranging from 44 to 51 mm, its capability to directly capture insects from water surfaces (Wilson and Mittermeier 2019) suggests a potential for piscivory. It is widely believed that the fishing behavior in bats evolved from their insectivorous habits, with the hypothesis that capturing insects from water represents an intermediate stage in this behavior (Aizpurua et al., 2013). Despite the relatively large hind feet of *M. horsfieldii*, exceeding half the length of the tibia, and their observed circular flight patterns above open water surfaces at a close distance of a few centimeters to search for insects, there is currently a lack of evidence for fishing behavior in *M. horsfieldii* (Aizpurua and Alberdi, 2018). Furthermore, their dietary composition has never been reported to date, leaving unanswered questions regarding whether fish forms part of its diet and what insect species they prey upon.

Hong Kong, situated at the southern coast of China, is the distribution range of *Myotis* species (Shek, 2006). It is a semi-island known for its diverse habitats, including various types of forests, coastal areas, wetlands, marine, and freshwater habitats, each hosting a wide array of arthropod and fish species (Dudgeon and Corlett, 2004). In this region, four *Myotis* species coexist, including *M. pilosus*, *M. horsfieldii*, *M. chinensis* (Chinese myotis), and *M. muricola* (whiskered myotis, locally rare). *Myotis chinensis* is bigger in size than *M. pilosus*, with a body length ranging from 91 to 97 mm (Wilson and Mittermeier 2019), and possesses relatively large hind feet compared to some other bat species. However, there are only a few studies conducted in China reporting its diets. Despite its large size, it is believed that *M. chinensis* exclusively feeds on insects. They are capable of preying on larger insects and capturing ground-dwelling insects through picking behavior, primarily consuming species in Coleoptera (e.g., carabid beetles), Orthoptera (grasshoppers), and Diptera (flies) according to an earlier study (Ma et al., 2008). Given the uncertainties surrounding the extent of piscivory by different *Myotis* species in the region, this study seeks to investigate the diets of *M. pilosus*, *M. horsfieldii*, and *M. chinensis* in Hong Kong by analyzing their fecal compositions using DNA metabarcoding.

Traditional microscopy for species identification of fecal or stomach contents presents several challenges. The identification of partially digested prey body parts under a microscope relies heavily on the expertise and experience of the analyst. Moreover, the process can be a time-consuming, subjective, and may have limitations in taxonomic resolution, potentially introducing biases or inconsistencies in prey identification (Svendsen et al., 2023, Massey et al., 2021). In contrast, DNA metabarcoding overcomes some of these challenges. It is a high-throughput technique that enables the analysis of a large number of samples and provides a higher taxonomic resolution even with small amounts of prey DNA (de Sousa et al., 2019, Monterroso et al., 2019).

By analyzing the prey identities and relative quantities of the three *Myotis* species, we can gain insights into the spectra of habitats bat individuals forage and prey they target in a habitat-diverse environment. Notably, both *M. chinensis* and *M. pilosus* are listed as Near Threatened species in the Red List of China’s Vertebrates (Jiang et al., 2016), with *M. pilosus* also being classified as a Vulnerable species in the IUCN Red List of Threatened Species (IUCN, 2023, Jiang et al., 2019). Hence, it would be important to further explore the dietary niche partitioning of these sympatric species. The findings would inform us their trophic relationships and what types of food resources and habitats are critical to their population sustainability, and significantly contribute to the conservation efforts for each of these species in the region.

In this study, we aim 1) to investigate the dietary compositions of the three *Myotis* species at both individual and species levels, 2) to estimate the dietary diversity within individuals and populations, 3) to assess the effect of environmental and host factors on the dietary compositions of these species, and 4) to determine the patterns of dietary niche partitioning among the *Myotis* species. We hypothesize that the three species might demonstrate dietary niche partitioning. We predict distinct dietary compositions for each species, resulting in minimal dietary overlap between species. Specifically, we predict that *M. pilosus* might include diverse fish taxa in its diet due to the diverse aquatic habitats close to the roosting sites. We also hypothesize that there is seasonal variation in the dietary compositions of the bat species. We predict a shift in food consumption pattern across different seasons.

## 2. Materials and Methods

### 2.1 Sample collection

Between 2018 and 2021, we visited water tunnels or caves in Hong Kong to capture *Myotis* bats during wet (April to September) and dry (October to March) season. We refrained from visiting roosting sites during the months when bats were overwintering or breeding/nursing, thus minimizing any potential disturbance. The locations include Lin Ma Hang (LMH01), Tai Lam Chung (TLC01 and TLC02), Pak Tam Chung (PTC), and Sai Kung (SK0, SK02, and SK03) (Fig. S1 and Table S1a-c) (Shek, 2004). We captured the bats using hand-held hoop nets and placed them in breathable sterilized bags. To prevent cross-contamination, each individual bat was kept in a separate sterilized bag. We identified the species and sex of each bat based on its morphology (Poon et al., 2023, Shek, 2006). The fresh feces from each bat were collected and placed in individual 2mL tubes. They were preserved in 100% ethanol and then kept at -80℃ until DNA extraction. In total, we have collected fecal samples from 62 *M. pilosus*, 51 *M. horsfieldii*, and 43 *M. chinensis*. Physical parameters of each bat, such as body weight and forearm length, were measured. After collecting samples and recording their physical parameters, all bats were released immediately back to the wild. Approvals for animal experiments were granted by the Department of Health (ref. 19-177 in DH/SHS/8/2/3 Pt. 30), the Committee on the Use of Live Animals (ref. 4963-19), and the Agriculture, Fisheries, and Conservation Department (AFCD; ref. 35 in AF GR CON 09/51 Pt.8).

### 2.2 DNA extraction and metabarcoding

The QIAamp Fast DNA Stool Mini Kit (Qiagen, Hilden, Germany) was used to extract the fecal DNA. We included negative controls and mock communities (Table S2a) during DNA extraction to later check for contamination during PCR. We quantified the fecal DNA using the Qubit dsDNA high-sensitivity (HS) assays on an Invitrogen Qubit 4 Fluorometer (Thermo Fisher Scientific, Massachusetts, US) (Huang et al., 2021).

All fecal DNA samples, mock communities (Table S2a), and negative controls were used for library preparation using three genetic markers through two-step PCR (Huang et al., 2022, Huang et al., 2021) (Supplementary Materials and Methods). The first pair of markers used was a universal pair (18s_SSU3_F: 5’GGTCTGTGATGCCCTTAGATG3’ and 18s_SSU3_R: 5’GGTGTGTACAAAGGGCAGGG3’) which targets the V7 region of 18S small subunit ribosomal DNA (rDNA; ca. 174bp) (McInnes et al., 2017). This pair of markers provides an overview of the dietary compositions of *Myotis* bats. To investigate whether fish was consumed by the three *Myotis* species, we used the mitochondrial 12S rDNA specific primers (12S_AcMDB07_HK_F: 5’GCCTATATACCRCCGTCG3’ and 12S_AcMDB07_ R: 5’GTACACTTACCATGTTACGACTT3’; ca. 283bp) to amplify fish DNA (Bylemans et al., 2018). Considering the important prey of *Myotis* bats, which consists of arthropods, we used a third pair of markers which amplifies the mitochondrial cytochrome c oxidase subunit I gene (COI; COI_Fwh2_F: 5’ GGDACWGGWTGAACWGTWTAYCCHCC3’ and COI_Fwh2_R: 5’ GTRATWGCHCCDGCTARWACWGG3’; ca. 219bp) (Vamos et al., 2017). This pair of markers offers a higher resolution in identifying the macroinvertebrate species consumed by the bats. For each marker, we generated a library multiplex by pooling libraries from different samples in an equimolar ratio. Subsequently, all three libraries were sequenced on an Illumina NovaSeq (PE 150bp) by Novogene (Hong Kong).

### 2.3 Sequencing data preprocessing

Raw paired-end DNA reads were merged by using the -fastq_mergepairs function in USEARCH v11.0.667 (Edgar, 2010). Primer sequences were removed with CUTADAPT v2.5 (Martin, 2011). The assessment of trimmed reads quality was completed with FastQC v0.11.9 (Wingett and Andrews, 2018) and VSEARCH v2.18.0 (Rognes et al., 2016). Only high-quality trimmed reads within the target lengths (18S: 130-180bp; COI: 130-210bp; 12S: 140-290bp) were retained for later analysis. These pre-processed reads were then dereplicated by using the - derep_fulllength command in VSEARCH. Chimeras and singletons were removed from the dereplicated reads by using USEARCH. Using the -usearch_global function in VSEARCH, all pre-processed reads were clustered into amplicon sequence variants (ASVs) based on 99% similarity.

Using the SINTAX algorithm in USEARCH, each ASV was assigned to the lowest identifiable taxonomic level with a confidential cutoff of 0.7 (Edgar, 2016). The ribosomal RNA database SILVA (Glöckner et al., 2017), the mitochondrial database MIDORI for eukaryotes (Leray et al., 2022), and MitoFish for fish (Zhu et al., 2023) were used as taxonomic classification reference databases for 18S, COI, and 12S sequences, respectively. We also assigned ASVs against the NCBI non-redundant nucleotide sequences database to obtain the best 1000 blast hits with a similarity higher than 99% and e-value lesser than 1e-50. The BASTA with the lowest common ancestor (LCA) algorithm (Kahlke and Ralph, 2019) helped us assign the lowest common taxonomic level shared by 80% of blast hits. To obtain a high taxonomic classification resolution, the results from SINTAX and LCA were combined by assigning ASVs with lower ranks of taxonomies.

Potential false-positive and contaminant ASVs were eliminated by comparing them with mock communities and negative controls (Table S2b-d). Non-diet ASVs (e.g., Fungi, Bacteria, and Algae) and unclassifiable ASVs (e.g., ASVs that could not be identified beyond the domain level) were discarded from the analysis. After ASVs cleaning, samples with reads number lower than 100 were subsequently excluded. To better present our taxonomic data in figures, we classified each identified taxon as high and low abundance taxon according to whether it occupies over 0.1% number of reads. High abundance taxa were grouped to the lowest taxonomic level, while low abundance taxa were grouped into higher taxonomic level. In this way, the taxa were classified into taxonomic categories (Table S3a-c) (Huang et al., 2022).

### 2.4 Data analysis

After data preprocessing, we used 18S data of 144 samples, 12S data of 48 samples, and COI data of 100 samples for downstream data analysis. In the 18S dataset, a total of 11 taxonomic categories were classified from 91 ASVs. The identifiable taxonomic levels categorized included three classes, eight orders, two families, and one genus (Table S3a). In the 12S dataset, 22 taxonomic categories were classified from 24 ASVs. The taxonomic levels included nine families and 12 species (Table S3b). In the COI dataset, 29 taxonomic categories were classified from 654 ASVs. The taxonomic levels included two classes, 18 orders, 13 families, 19 genera, and 18 species (Table S3c). In the statistical analysis, we calculated (1) the percentage of read count for each taxon in a sample (relative read abundance, RRA), (2) the percentage of occurrence for each taxon in a sample (weighted percentage of occurrence, wPOO), and (3) the proportion of samples in which a taxon is detected (frequency of occurrence, FOO). The RRA or wPOO at the population level is presented as the mean of RRA or wPOO of all individual samples of a bat species (Lee et al., 2021). We conducted these analysis in R v4.2.1 and visualized with the R package ggplot2 v3.4.2 (Hadley, 2016).

#### 2.4.1 Diet diversity analysis

To determine the relationships between sample numbers and dietary species Chao2 diversity, we generated rarefaction curves using hill numbers from ASVs or taxa (*q*=0) via the R package iNext (Hsieh et al., 2016) (Fig. S2). Using hill numbers based on variant *q* values (the order of diversity), we estimated the diet diversity of three *Myotis* species at both individual (alpha diversity) and population/species (gamma diversity) levels. Hill number only considers the occurrence of each ASV when *q*=0. The weight of species abundance in hill number increases when *q* value increases. The hill number is equivalent to the exponential of Shannon’s diversity index and inversion of Simpson’s dominance at *q*=1 and *q*=2, respectively (Alberdi and Gilbert, 2019). Pairwise diversity comparisons at the individual level (alpha) between *Myotis* species were carried out using the Kruskal‒Wallis test, followed by Benjamini-Hochberg correction (p<0.05).

Beta diversity refers to the differences in dietary taxa compositions between individual samples. To analyze beta diversity, we calculated pairwise binary Jaccard dissimilarity distances and pairwise Bray‒Curtis dissimilarity distances using the occurrence of each ASV and the fourth root transformed RRA of each ASV, respectively. We visualized the dissimilarities with principal coordinates analysis (PCoA) with the plot_ordination function in R package phyloseq v1.42.0 (McMurdie and Holmes, 2019), and fitted the effect of each taxonomic category into the PCoA result using the envfit and ordiArrowMul function in R package vegan v2.6.4 (Oksanen et al., 2022). Pairwise permutational multivariate analysis of variance (PERMANOVA) tests were performed to investigate the separation of dietary compositions between the *Myotis* species. Significant interspecific composition variants observed in PERMANOVA tests are premised on the intraspecific homogeneity, which were tested by betadisper function. We also carried out similarity percentage (SIMPER) analyses to assess the contribution of each taxon to the difference between *Myotis* species by using R scripts simper_pretty.R (Steinberger et al., 2016), which were then checked by the Kruskal-Wallis test by using R scripts R_krusk.R (Steinberger, 2020).

Within each species, we performed multiple analyses to determine the contribution of host variables (i.e., sex) and environmental variables (i.e., sampling seasons and sampling locations) to the diversity variation. Data from SK01-03 and PTC were combined for analyses due to their close geographic proximity. Generalized linear models were performed to evaluate the effect of variables on alpha dietary diversity by using the logarithmic hill number of order *q*=1 as the dependent variable. We visualized the beta diversity dissimilarities of variable groups using PCoA based on Bray-Curtis and Jaccard dissimilarity distances with fitted taxonomic categories. PERMANOVA and corresponding beta-dispersion tests were used to assess the beta diversity variations among variable groups. We identified taxa that significantly contributed to the composition variation among variable groups, using the SIMPER test, which was checked by the Kruskal‒Wallis test. The dietary composition overlap between bat individuals was measured by Pianka’s niche overlap index using the R package “spaa” v0.2.2 (Zhang et al., 2016). A network was generated to visualize individual-level diet overlap by using the “qgraph” package v1.9.5 (Pedersen et al., 2017).

## 3. Results

### 3.1 Dietary compositions of the three *Myotis* bat species

Based on 18S data, the three *Myotis* species primarily preyed upon insects, arachnids, and/or ray-finned fish (Actinopteri) (Fig. 1 and S3). Specifically, flies (dipterans), moths and/or butterflies (lepidopterans), orthopterans, and true bugs (hemipterans), were the most commonly consumed insect groups. Arachnids such as spiders (Araneae) and mites (Mesostigmata), as well as fish also formed part of their diet.

**Fig. 1.**
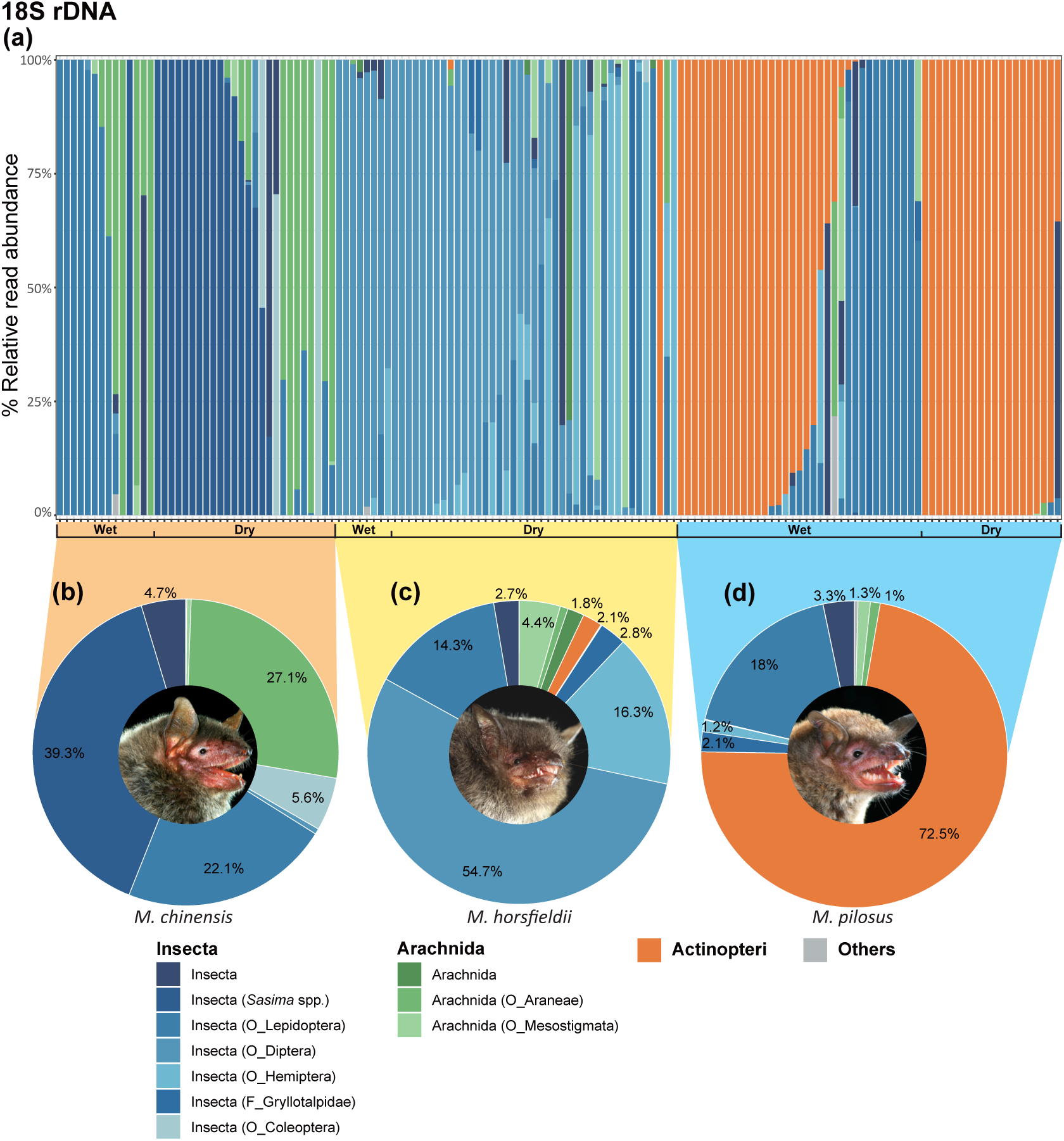
Dietary composition of *Myotis* bat species based on taxa detected in fecal samples using the 18S rDNA marker. The relative read abundance of each taxon were visualized at both (a) individual level (n=144) and (b-d) species level for (b) *M. chinensis* (n=40), (c) *M. horsfieldii* (n=49), and (d) *M. pilosus* (n=55). Prey in classes with read abundance lower than 0.1% of all taxa detected were grouped into “Others”. Only taxa with abundance higher than 1% in relative read abundance are shown. F, Family. O, Order. Bat photos © Agriculture, Fisheries and Conservation Department.

The three *Myotis* species exhibited noticeable interspecific variations in their primary food sources (Fig. 1b-d). *Myotis chinensis* (18S, n=40) primarily consumed insects (72% RRA), particularly bush crickets (*Sasima* spp. in order Orthoptera at 39%) and moths and/or butterflies (22%). Additionally, spiders accounted for 27% of their diet (Fig. 1b and Table S4). *Myotis horsfieldii* (18S, n=49) mainly hunted smaller size insects (91%), with flies (55%) and true bugs (16%) being the primary prey. They consumed arachnids (7%) but in smaller proportions (Fig. 1c; Table S4).

Fish was mainly found in the diets of *M. pilosus* (n=55; Fig. 1d; Table S4). The 18S data also revealed the presence of fish contents in two samples of *M. horsfieldii*, which were not detected by the 12S (Fig. 1a and 1c; Table S5a-b). To validate the presence of fish in the samples of *M. horsfieldii*, we first confirmed the bat species identity by DNA barcoding using primers SFF_145f and SFF_351r (Walker et al., 2016), and then we confirmed the fish contents of these samples by performing PCR using the 18S and 12S primers (section 2.2) and Sanger sequencing. We concluded that the results were consistent with the DNA metabarcoding finding that fish was identified in these two samples.

*Myotis pilosus* (18S, n=55) displayed significant intraspecific differences in diets (Fig. 1 and 2a; Table 6a-b). Approximately one-fifth (9/55) of the individuals were found to exclusively feed on macroinvertebrates, with a majority of these individuals found in the wet season. These individuals primarily prey on moths and/or butterflies. In contrast, the remaining individuals preyed on fish in varying degrees, constituting 6-100% of their individual diets (Fig. 1a and 1d; Table S5a-b).

**Fig. 2.**
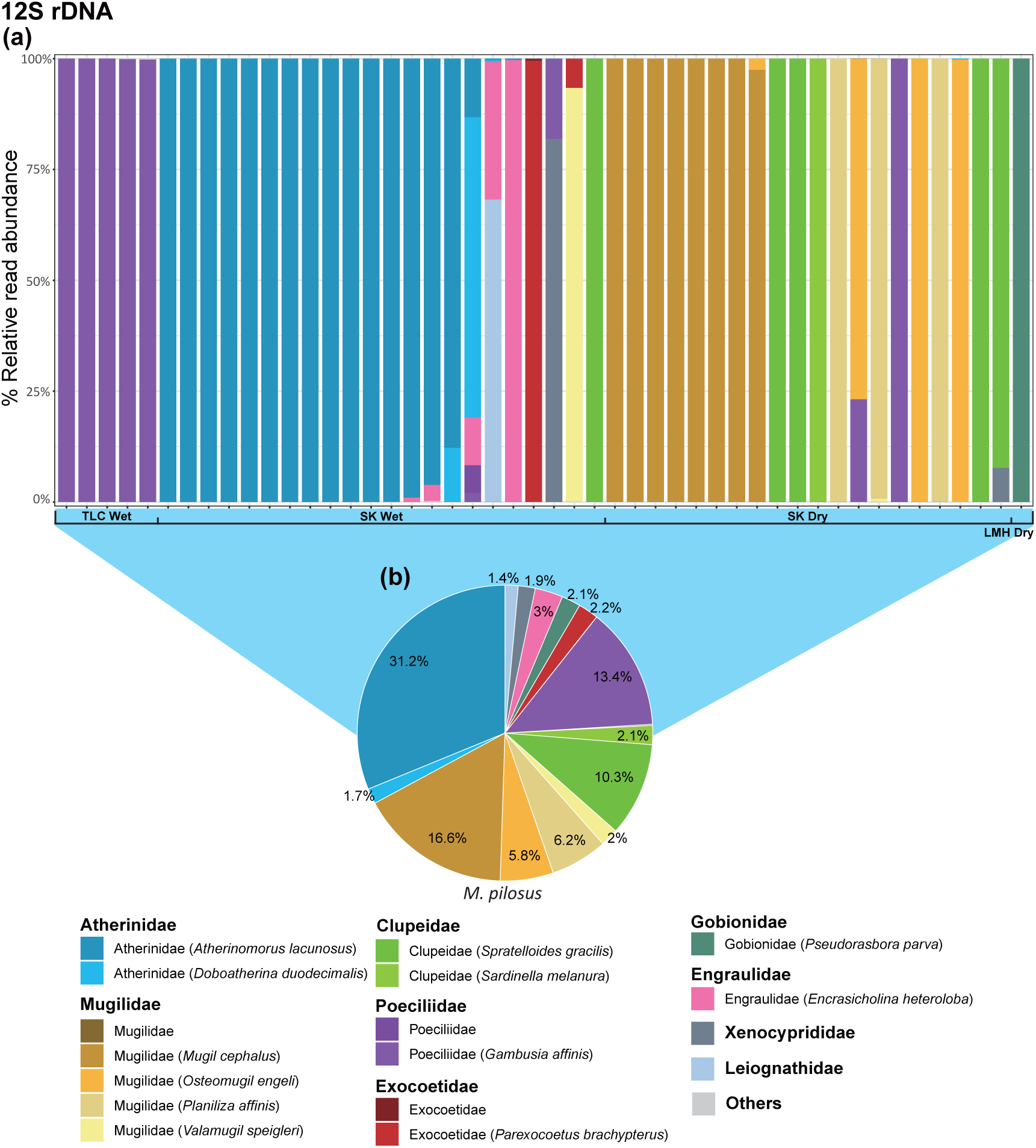
Dietary composition of *Myotis pilosus* based on taxa detected in fecal samples using the 12S rDNA marker. The relative read abundance of each taxon were visualized at both (a) individual (n=48) and (b) species levels. Individual bars were grouped by sampling locations and seasons. Prey in classes with read abundance lower than 0.1% of all taxa detected were grouped into “Others”. Only taxa with abundance higher than 1% in relative read abundance are shown. Refer to Fig. S1 for locations. SK, Sai Kung (including Pak Tam Chung); TLC, Tai Lam Chung; LMH, Lin Ma Hang.

Based on the 12S metabarcoding data, all individuals that consumed fish were *M. pilosus* (Fig. 2a and S4; Table S6a-c). Both freshwater, marine, or brackish fish were found in the diet of *M. pilosus*, at least twelve fish species were detected. *Myotis pilosus* primarily hunted marine fish, with Old World silversides (Atherinidae) making up 33% of their diet, including wide-banded hardyhead silverside (*Atherinomorus lacunosus*, max. size <25cm) at 31%. Mullets (Mugilidae) comprised 31% of their diet, with flathead grey mullet (*Mugil cephalus*, common max. size <50cm) making up 17%, and eastern keelback mullet (*Planiliza affinis*, max. size <40cm) and kanda mullet (*Osteomugil engeli*, max. size <48cm) each contributing about 6%. Herrings and sprats (Clupeidae) accounted for 12% of their diet, with silver-stripe round herring (*Spratelloides gracilis*, max. size <11cm) being commonly consumed at 10%. Mosquitofish (*Gambusia affinis*, max. size <5.1cm, from the family Poeciliidae) was the dominant freshwater fish at 13% (Froese and Pauly, 2024). Notably, mosquitofish was mainly consumed by individuals from Tai Lam Chung (Fig. 2; Table S6a-b).

The COI data offered a more detailed revelation of the macroinvertebrate prey consumed by the *Myotis* species (Fig. 3 and S5; Table S7a-c). Overall, the majority of prey consisted of dipterans (about 32%), such as lake flies in the family Chironomidae (e.g., *Procladius culiciformis*, *Glyptotendipes tokunagai*, *Chironomus flaviplumus*, etc.); lepidopterans (14%), including grass moths like *Syntonarcha iriastis* and *Cirrhochrista brizoalis*; orthopterans (13%), including crickets like *Mecopoda* spp. (bush crickets) and *Gryllotalpa* spp. (mole cricket); and spiders (12%), such as northern golden orb weaver (*Nephila pilipes*) (Fig. 3; Table S7a).

**Fig. 3.**
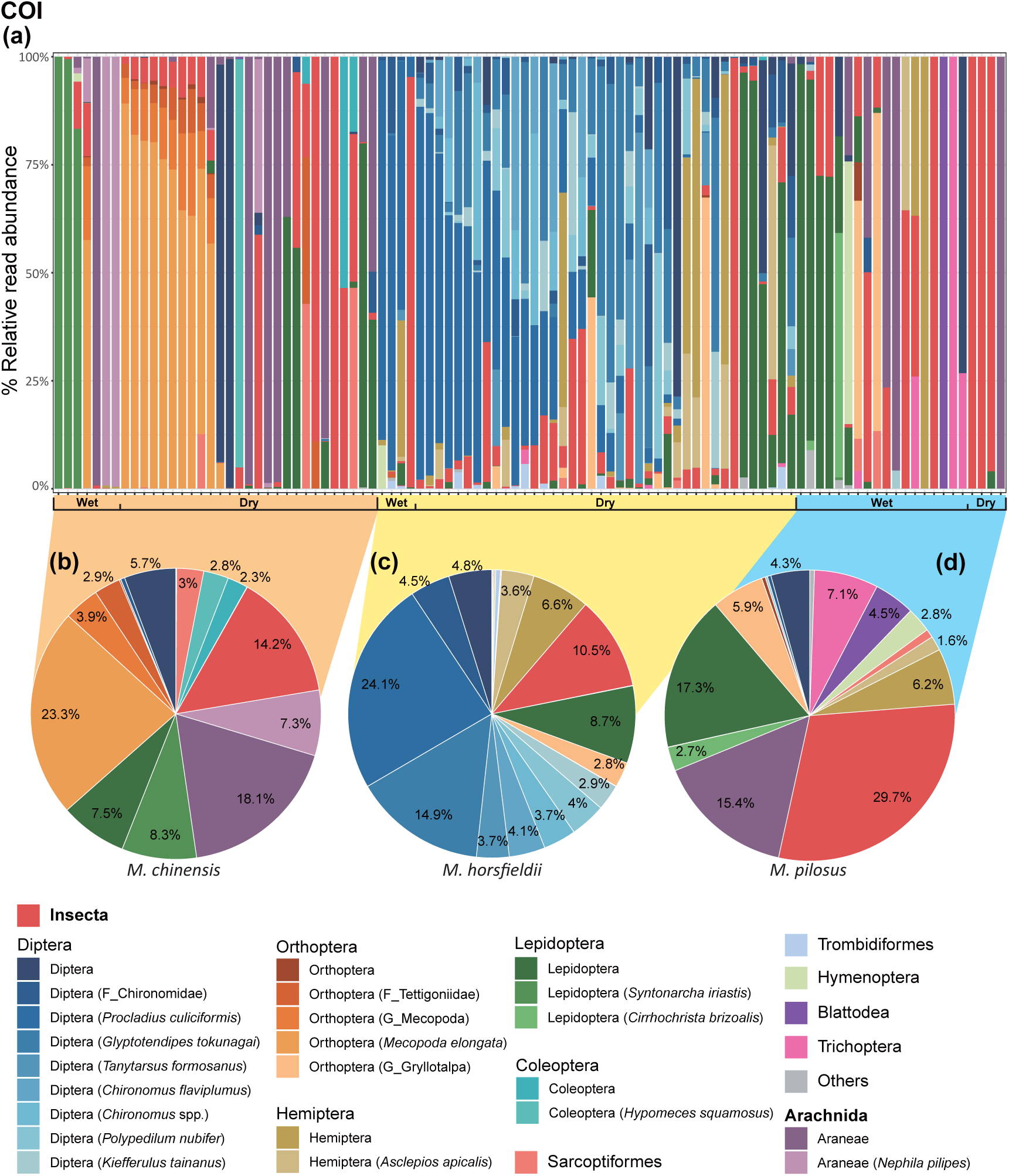
Dietary composition of *Myotis* bat species based on taxa detected in fecal samples using the COI marker. The relative read abundance of each taxon was visualized at both (a) individual level (n=102) and (b-d) species level for (b) *M. chinensis* (n=34), (c) *M. horsfieldii* (n=44), and (d) *M. pilosus* (n=22). Prey in classes with read abundance lower than 0.1% of all taxa detected were grouped into “Others”. Only taxa with abundance higher than 1% in relative read abundance are shown. G, Genus. F, Family. O, Order.

The macroinvertebrate orders in diets revealed by COI were consistent with those unveiled by 18S. Notable variations were observed in the dietary compositions among *Myotis* species, as revealed by COI. *Myotis chinensis* (COI, n=34) primarily preyed on arthropods in several orders, such as bush cricket *Mecopoda elongata* (23%) from Orthoptera (30%), *S. iriastis* (8%) from Lepidoptera (16%), and *N. pilipes* (7.5%) from Araneae (25%). Spiders were found in more than 50% of the *M. chinensis* samples (Fig. S5b; Table S7a). In *M. horsfieldii* (COI, n=44), smaller size insects like dipterans accounted for more than 66% of their total diet, with *P. culiciformis* (24%) and *G. tokunagai* (15%) being the main contributors. More than 50% of the *M. horsfieldii* samples contained various lake fly species. Lepidopterans and hemipterans, on the other hand, made up 9% and 10% respectively (Fig. 3c; Table S7a). Although a larger proportion of 30% of insects remained unidentified compared to the other two bat species, the primary prey of *M. pilosus* (COI, n=22) consisted of arthropods from various orders, including Lepidoptera (20%), Araneae (15%), Orthoptera (7%), Hemiptera (8%), and Trichoptera (caddisflies, 7%) (Fig. 3d; Table S7a).

### 3.2 Alpha and gamma diversity of consumed taxa

The alpha diversity of individual diets indicated that the diets of *M. horsfieldii* exhibits the highest diversity compared to the other two *Myotis* species, as shown by 18S and COI data (Fig. 4 and 5; Table S8a-b). Notably, there is a distinct decrease in hill numbers as the *q* value increases, particularly observed in the macroinvertebrate (COI) composition of *M. horsfieldii* and *M. chinensis* diets (Fig. 5; Table S8b). This suggests that although these bats consume a greater variety of macroinvertebrates compared to *M. pilosus*, their prey compositions are highly uneven, with certain taxa dominating their diets. Furthermore, the macroinvertebrate (COI) compositions of *M. chinensis* were more unevenly distributed compared to *M. horsfieldii* (Fig. 5; Table S8b). In the case of individuals of *M. pilosus* that consumed fish, mostly two to three fish species were detected in their diets (12S data; Fig. 2a; Fig. 5). A similar pattern of prey consumption based on 18S data was also observed at the population level as reflected by gamma diversity, with the dietary composition of *M. horsfieldii* being the most diverse (Fig. 5; Table S8a). Despite having the lowest macroinvertebrate (COI) diversity in their diets, *M. pilosus* displayed comparable diversity of overall (18S) diets to *M. chinensis* at the population level (Fig. 5; Table S8a-b).

**Fig. 4.**
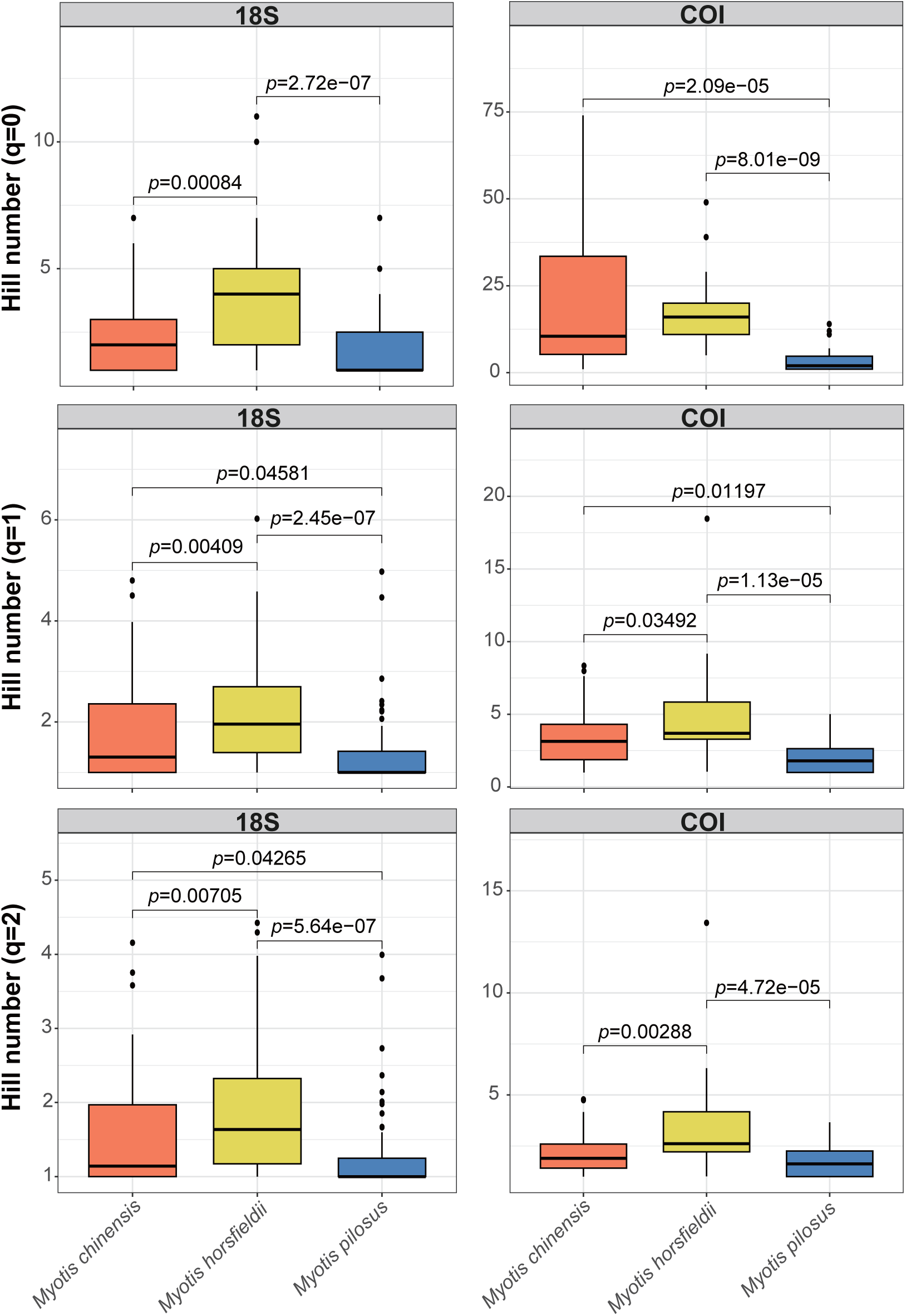
Pairwise alpha diversity comparison between *Myotis* species. Alpha diversity is represented by using the hill numbers of order *q*=0 (upper panel), *q*=1 (middle panel), and *q*=2 (lower panel), which were calculated based on the ASVs abundance detected by18S rDNA (left panel) and COI (right panel) marker at individual level. Only the *p*-values of species pairs which have significant different alpha diversity were shown (*p*<0.05).

**Fig. 5.**
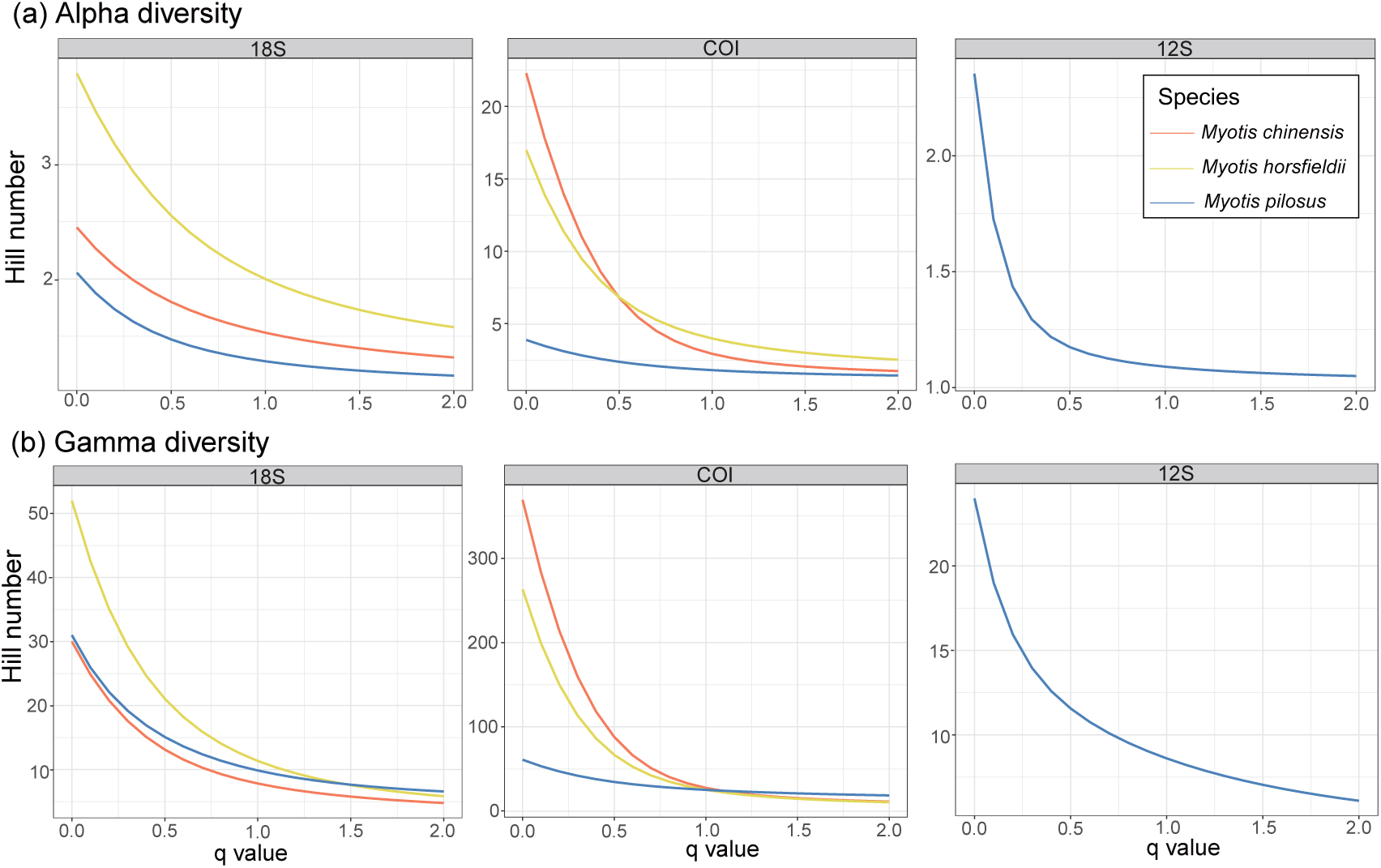
Dietary diversity for each *Myotis* represented in the hill numbers of variant order *q*. The hill numbers were calculated at individual level and species level to estimate the (a) alpha diversity and (b) gamma diversity, respectively. The ASVs abundance of each sample were detected by using 18S rDNA (left panel), COI (middle panel), and 12S rDNA (right panel) marker gene.

### 3.3 Effects of environmental and host factors on the dietary compositions

According to the PERMANOVA tests conducted using Bray‒Curtis and Jaccard dissimilarity distances, the diets of *M. pilosus* were found to differ significantly between seasons and locations based on 12S (Table S9a-b) and 18S (Table S10a-b) data. Specifically, during the wet season, there was a higher consumption of macroinvertebrates by *M. pilosus*, mainly lepidopterans as evidenced by the 18S data, while in the dry season more fish was consumed. Moreover, the fish compositions of *M. pilosus* from SK, where a majority of samples that were collected from, exhibited notable variations between the wet and dry seasons (Fig. 2a). The SIMPER analysis results further identified the main contributors to location difference. Old World silverside (Atherinidae) were consumed more at SK, while mosquitofish were consumed more at TLC (Fig. 6a-b and Table S11a-b) (Froese and Pauly, 2024).

**Fig. 6.**
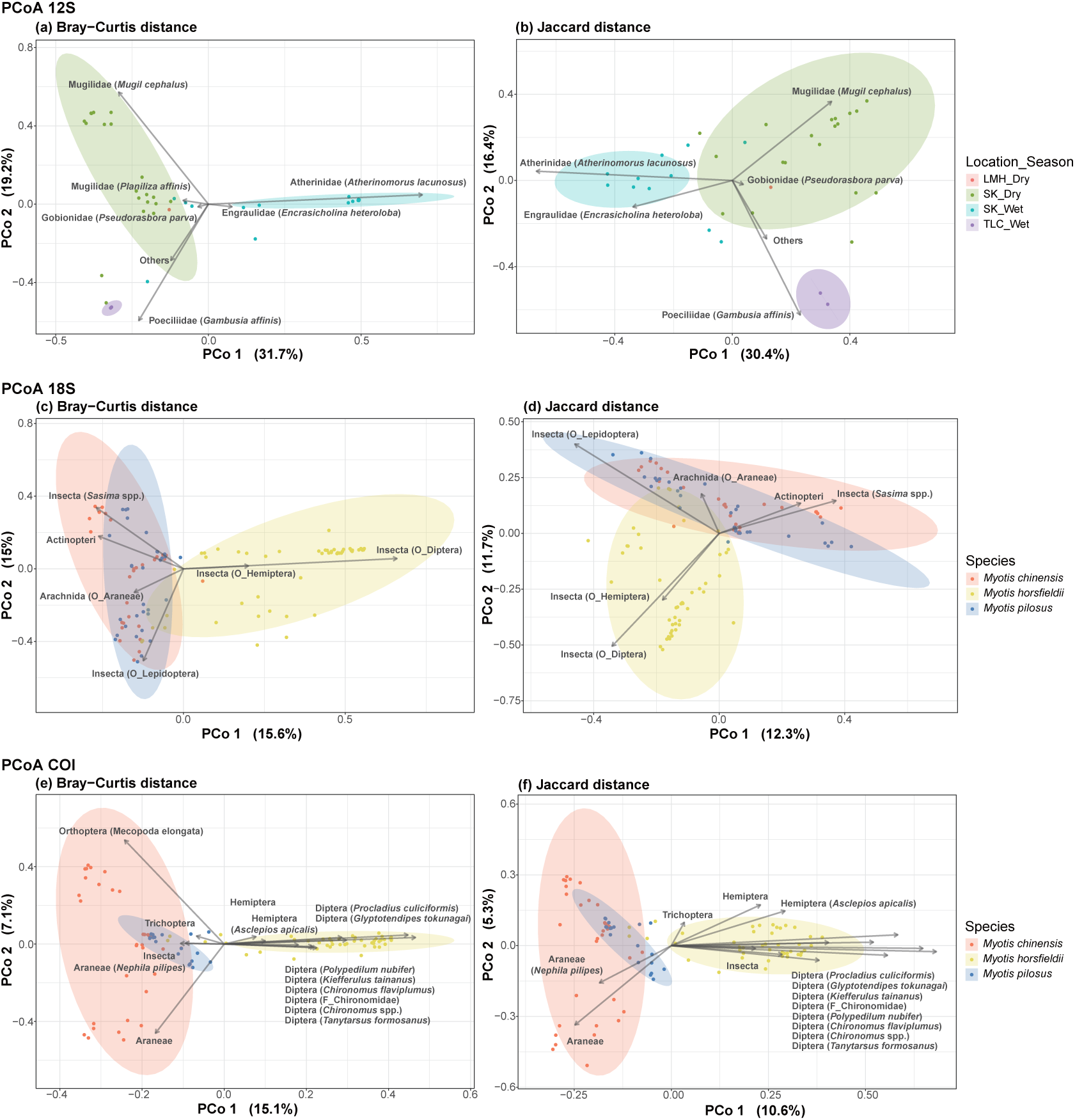
Principal coordinate analysis (PCoA) of dietary compositions in (a, b) *Myotis pilosus* and (c-f) three *Myotis* species based on (a, c, e) the pairwise Bray‒Curtis dissimilarity distances calculated from the fourth root transformed RRA of each ASV, and (b, d, f) the pairwise Jaccard dissimilarity distances calculated from the occurrence of each ASV. Each point corresponds to one sample, with the color indicates (a, b) sampling locations, seasons, and (c-f) *Myotis* species. The ASVs were identified by the (a, b) 12S rDNA, (c, d) 18S rDNA, and (e, f) COI marker gene. Taxonomic categories significantly contributing to the difference between groups were shown (see Table S11, S15, and S16). Refer to Fig. S1 for locations. SK, Sai Kung (including Pak Tam Chung); TLC, Tai Lam Chung; LMH, Lin Ma Hang.

Similarly, the diets of *M. chinensis* also differed significantly between the two seasons based on both 18S (Fig. 1a and Table S10a-b) and COI (Fig. 3a and Table S12a-b) data. The 18S data showed that during the wet season, *M. chinensis* mainly consumed moths and/or butterflies (Lepidoptera), while in the dry season, they consumed a large proportion of *Sasima* bush crickets (Orthoptera) (Fig. 1a). Although both GLM and PERMANOVA analyses consistently demonstrated a significant divergence in the seasonal diets of *M. horsfieldii* (Table S12a-b and S13), it’s important to note that the small sample size (n=4) from the wet season in the COI dataset may limit the accuracy of these findings. Further investigations are required in the future to confirm this result. Furthermore, the PERMANOVA analysis revealed an additional noteworthy disparity in the diets of the two sexes of *M. horsfieldii* (Table S12a-b), which was mainly attributed to the distinct consumption of dipterans, lepidopterans, and hemipterans by each sex (Table S14a-d).

### 3.4 Dietary niche partitioning between the three *Myotis* species

Based on the PCoA analysis, 18S and COI data revealed highly similar patterns of dietary niche partitioning among the three *Myotis* species (Fig. 6c-f). The PCoA analysis using Bray‒Curtis and Jaccard distances within each of the 18S and COI dataset also revealed consistent patterns. The overall (18S) and macroinvertebrate (COI) diets of *M. horsfieldii* were distinctly differentiated from those of *M. pilosus* and *M. chinensis*, while the dietary compositions of *M. pilosus* and *M. chinensis* were more overlapping with each other (Fig. 6c-f). The SIMPER analysis results indicated the main contributors to this differentiation in *M. horsfieldii* were predominantly dipterans and hemipterans, which constituted a significant portion of their diet (Fig. 6c-d; Table S15a-b). Specifically, the true bugs included water striders (e.g., *Asclepios apicalis*) while dipterans included a group of lake flies (e.g., *P. culiciformis*, *G. tokunagai*, *Kiefferulus tainanus*, *Polypedilum nubifer*, *Chironomus* spp. etc.) (Fig. 6e-f; Table S16a-b).

*Myotis chinensis* also differentiated from *M. horsfieldii* and *M. pilosus* by consuming a considerable proportion of bush crickets (Orthoptera), such as *Sasima* spp. and *M. elongata*, as well as spiders (Araneae, including *N. pilipes*) (Fig. 6c-f; Table S15a-b and S16a-b). On the other hand, the dietary niche of *M. pilosus* differed from the other two species by including a large portion of fish (Actinoptera) in their diets (Fig. 6c-d and Table S15a-b). *M. pilosus* also differentiated from *M. chinensis* by consuming caddisflies (Trichoptera) (Fig. 6e-f; Table S16a-b).

At the individual level, *M. horsfieldii* showed greater intraspecific overlap of individual diets compared to those of *M. chinensis* and *M. pilosus* (Fig. S6). This suggests the dietary compositions among *M. horsfieldii* individuals were more similar, and the dietary compositions among individuals of *M. chinensis* and *M. pilosus* were more variable.

## 4. Discussion

### Fish eating behavior of *Myotis*

This study provides new insights into the dietary compositions of *M. pilosus* and *M. chinensis* in coastal habitats, expanding on previous research that primarily focused on inland regions of China. Additionally, our research contributes to the understanding of the dietary compositions of *M. horsfieldii*, a species found in southeast and south Asia that has not been previously studied in terms of its foraging ecology. Contrary to earlier studies conducted in inland regions of China, which suggested that *M. pilosus* primarily fed on cyprinids in freshwater habitats year-round (Ma et al., 2006, Chang et al., 2019), our research in Hong Kong unveiled a different dietary pattern. Our findings show that *M. pilosus* in Hong Kong did not prey on cyprinids but instead exhibited a diverse diet, targeting a wide variety of fish prey species from nine different families. These prey species encompassed marine, brackish, and freshwater fish, highlighting the versatility of *M. pilosus* in its feeding habits. Moreover, we observed variations in the species compositions consumed by *M. pilosus* across different seasons. For instance, marine species such as *A. lacunosus*, *Doboatherina duodecimalis* (tropical silverside, max. size 11cm), *Parexocoetus brachypterus* (sailfin flying fish, max. size 13cm), and *E. heteroloba* were only fished by *M. pilosus* during the wet season at Sai Kung. This was observed specifically at sampling site SK01 but not the other sites. Worth noting is that SK01 is located 1.7 km or less away from the nearest shore at the southwest, making it a likely site for *M. pilosus* to fish rather than flying north-or eastward to reach the shore, which is located over 3 km away.

*Atherinomorus lacunosus*, which forms large schools along sandy shorelines, serves as an important food source for *M. pilosus*. This fish species forages nocturnally to capture zooplankton that migrate vertically to the upper water column during nocturnal hours (Skibinski, 2005). This feeding behavior creates large amount of ripples that trawling bats like *M. pilosus* can detect. Another noteworthy prey species is *P. brachypterus*. Sailfin flying fish is commonly found in coastal waters in large shoals and possessing elongated, wing-like pectoral fins, which enable it to leap out of the water and glide rapidly for considerable distances above the surface, an adaptation to evade underwater predators. During the breeding season, spawning flying fish aggregate in abundance near the surface at night, with many vigorously jumping and flying out of the water to release ripe eggs and sperm (Stevens et al., 2003, Digo et al., 2015, Lewis, 1961), exposing them to attack by aerial predators, such as *M. pilosus*.

During the dry season in Sai Kung, *M. pilosus* captured the marine fish species *Spratelloides gracilis* and *Sardinella melanura* (blacktip sardinella, common size 10cm). Clupeids, such as *S. gracilis* and *S. melanura*, are primarily forage fish known for their high egg production. For instance, *S. gracilis* spawns near the water surface, releasing approximately 1600 eggs per unit body mass (Dalzell and Wankowski, 1980). Although previous studies in China have reported crustaceans as prey for *M. pilosus* and other bat species (Wang et al., 2024), our study did not identify any crustaceans in the diet of *M. pilosus*. Based on our results, one notable target species for *M. pilosus* is *M. cephalus*, along with other mullet species, during dry months. *Mugil cephalus* is known for congregating in schools over sand or mud in coastal and brackish waters. In Hong Kong, this species exhibits two short spawning peaks in winter (dry) months, resulting in a large numbers of juveniles appearing inshore, especially in estuaries, during February and March (Sadovy and Cornish, 2000).

In the Tai Lam Chung woodland area, which is an inland, *M. pilosus* primarily preys on the exotic freshwater poeciliid, *G. affinis*. The southern shore is obstructed by highland with an elevation of at least 400m, and it is at least 9 km away from the nearest shore in any direction (Fig. S1). *Gambusia affinis* is a freshwater species and a widespread non-native fish in Hong Kong, originating from North America and released by the Hong Kong Government for mosquito control purposes around 1940 (Dudgeon and Corlett, 2004). Surveys on the distribution of *G. affinis* in Hong Kong have reported its presence in the lower reaches of the closest streams within less than 1km of TLC forest (Tsang and Dudgeon, 2021a). Experiments conducted on the ecological effects of *G. affinis* have shown that it reduces the abundance and richness of invertebrates and alters assemblage compositions in Hong Kong wetland (Tsang and Dudgeon, 2021b). Our findings suggest that *M. pilosus* may play a potential role in controlling invasive poeciliids in the local ecosystem. Similar findings have been reported in *M. capaccinii* in northwest Israel and eastern Iberian Peninsula, where *M. capaccinii* fed on exotic *Gambusia* spp. (Aizpurua et al., 2014, Levin et al., 2006). Mosquito fish frequently swim close to water surface, using their upturned mouth to break the surface and capture floating insects. Furthermore, the decrease in oxygen during the night may compel these fish to come to the water surface for breathing (Aizpurua et al., 2014, Levin et al., 2006).

Our study has made an intriguing discovery regarding the feeding habits of *M. horsfieldii*. We reveal two individuals *M. horsfieldii* from Sai Kung have included fish in their diet. Interestingly, we have found the first case of *M. horsfieldii* preyed on *Asclepios apicalis*, a water strider species that is also targeted by *M. pilosus*. While most water striders are known for their capacity to glide effortlessly on the calm freshwater surface, *Asclepios* spp. are considered as sea skaters which are found primarily in brackish water along coasts (Poolprasert et al., 2022, Andersen and Foster, 1992). *Myotis horsfieldii* might be drawn to this food source due to the consistent ripples produced by *Asclepios* as they manoeuvre across water’s surface. Reports on *M. horsfieldii*’s foraging behavior are limited, but one literature mentions that *M. horsfieldii* typically flies in circles around 10 cm above water to skim insects, and they roost near water sources (Wilson and Mittermeier 2019). Our finding suggests that *M. horsfieldii* may capture fish while foraging insects over waters. Despite this finding, it should be noted that fish was only detected in very few individuals of *M. horsfieldii*. Therefore, further investigation is needed to determine whether *M. horsfieldii* actively engage in fishing or if those individuals simply mistakenly catch live or dead fish on the water surface while foraging for insects. Our discovery that *M. horsfieldii*, which was not previously known to consume fish, does in fact include fish in their diets has significant implications for the evolution of fishing behavior. The active foraging of aquatic insects by bats may lead to occasional consumption of fish, which in turn creates the selective pressure that drives the evolution of fishing behavior.

### Foraging niches of the three *Myotis* species

We observed a higher proportion of fish in the diet of *M. pilosus* during the dry season compared to the wet season, with a significant increase in insect components during the wet season, which is consistent with Ma et. al. (2006) but different from Chang et al. (2018). Our results showed that this increase in insect consumption was due to a 43% rise in the number of individuals preyed on macroinvertebrates during the wet season (22/35) compared to the dry season (4/20). Furthermore, while there were some variations in the insect dietary compositions compared to previous research, our study revealed a similar finding to Chang et al. (2019) in terms of *M. pilosus* primarily preying on lepidopterans in the wet season. Based on the nocturnal behavior of most moths and the fact that the lepidopterans identified in our study are all moth species, such as *S. iriastis* and *C. brizoalis* consumed by *M. pilosus*, we suggest that most of the lepidopterans preyed upon by *M. pilosus* are moths. Surveys on the abundance of forest invertebrates in Hong Kong have shown that local lepidopterans vary seasonally, with both number and biomass experiencing winter (dry season) lows and early summer (in May during wet season) maxima (Kai and Corlett, 2002). During the wet season, local *M. pilosus* also targeted trichopterans. Trichopterans are moth-like and closely related to lepidopterans, which are nocturnal and commonly associated with freshwater bodies (De Moor and Ivanov, 2008).

Here, we provide additional evidence for the specialization of *M. pilosus* in fishing, as we have consistently found fish in its diet throughout the year. It is worth noting that the abundance of lepidopterans in local terrestrial habitat is higher during summer (Kai and Corlett, 2002), this high abundance likely explains why some individuals of *M. pilosus* have shifted their target prey from fish to insects during this time. This shift may be driven by higher energetic profit associated with capturing a larger number of moths compared to capturing fish. Despite the higher nutritional value of a single fish compared to a single moth, catching fish likely incurs a greater energy cost (Aizpurua et al., 2013). This is due to the heavier body mass of fish and the increased effort required to accurately detect fish underwater and subsequently pull it out of the water (Aihartza et al., 2008). This results in lower capture efficiency when targeting fish (Altenbach, 1989). Therefore, our study highlights the adaptive feeding behavior of *M. pilosus*, which adjusts its diet based on prey availability, underscoring the significance of this feeding strategy.

Our results show that there is a greater overlap in the macroinvertebrate diet between *M. pilosus* and *M. chinensis*, while the diet of *M. horsfieldii* is distinct from the other two species. We found that local *M. chinensis*, similar to *M. pilosus*, also hunted a higher proportion of lepidopterans, such as grass moth, *S. iriastis*, during the wet season. Another noteworthy foraging habit shared between *M. chinensis* and *M. pilosus* is their considerable consumption of non-flying nocturnal arthropods, specifically spiders like the northern golden orb weaver (*N. pilipes*). *Nephila pilipes* typically build intricate webs to capture prey on bushes and trees near water sources (Harvey et al., 2007). Spiders have been documented as part of the diet of several other *Myotis* species, such as *M. myotis*, *M. emarginatus*, *M. lucifugus*, *M. nattereri*, *M. evotis*, and *M. septentrionalis* (Goiti et al., 2011, Maucieri and Barclay, 2021, Swift and Racey, 2002, Kaupas and Barclay, 2018). However, previous studies did not report spiders as a food source for *Myotis* bats in China (Ma et al., 2003, Ma et al., 2008, Ma et al., 2006). Moreover, *M. chinensis* mainly preyed on orthopterans, especially bush crickets, such as *Mecopoda* and *Sasima* spp., during the dry season. The consumption of spiders and orthopterans suggests that *M. chinensis* is capable of foraging in cluttered environments of woodlands and gleans arthropods from various substrate surfaces, such as spider webs and/or foliage in bushes. While there are some geographical variations in dietary composition, our findings agree with Ma et al. (2008), which remarked that conspecifics in Beijing prey on ‘non-wing beating’ insects such as coleopterans (e.g., ground beetles in Carabidae) and orthopterans while lacking hymenopterans (sawflies, wasps, bees, and ants) and lepidopterans (Ma et al., 2008).

Based on our results, we found that the diet of *M. horsfieldii* is distinctive, which prominently includes a high consumption of smaller flying insects, specifically dipterans. Notably, *M. horsfieldii* preys on a wide variety of lake fly species from the Chironomidae family, leading to a higher dietary diversity compared to *M. chinensis* and *M. pilosus*. Conversely, our observations indicate that spiders, orthopterans, and coleopterans were seldom found in their excrement. In the forests of Hong Kong, dipterans constitute the majority of the insect biomass, and their abundance did not exhibit seasonal variation (Kai and Corlett, 2002). Numerous chironomid species bear a striking resemblance to mosquitoes and adults swarms in terrestrial habitats serve as vital sources of food for bats (Puig-Montserrat et al., 2020, Beck, 1995). Local *M. horsfieldii* also targets hemipterans and lepidopterans, though to a lesser extent compared to dipterans.

Our findings indicate that *M. pilous* is primarily a fish-eating bat, while *M. chinensis* and *M. horsfieldii* primarily feed on insects. One plausible reason for this dietary differentiation pattern could be differences in their morphological characteristics. Previous study suggested that the body size of aerial hunting bats limits their weight-carrying capacity, and fishing bat species tend to have relatively larger body sizes and longer hind feet that give them morphological advantages to fish hunting (Chang et al., 2019). Despite *M. chinensis* (91-97mm body length, 25-30g) (Wilson and Mittermeier 2019) and *M. pilosus* (51-65mm, 11.7-32.5g) exhibiting larger body sizes and heavier body masses, *M. pilosus* share a similar hind foot to forearm length ratio (0.31±0.014) to other fishing bats (0.31±0.051) (Chang et al., 2019), while the ratio for *M. chinensis* (0.26) is more similar to those of other non-fishing bats (0.25±0.038). *Myotis horsfieldii* (44-51mm, 5.0-7.5g) (Wilson and Mittermeier 2019) is smaller in size compared to the other two species. It has a low value ratio (0.23), which is also more akin to other non-fishing trawling bats (Chang et al., 2019). Apart from the influence of morphological adaptations on prey capturing ability, other factors contributing to the dietary differentiation pattern could include differences in prey detection ability through variations in echolocation call structures during foraging (Aizpurua and Alberdi, 2018).

This dietary study uncovered the overlapping macroinvertebrate diets of *M. pilosus* and *M. chinensis*, as well as the distinct diet of *M. horsfieldii*, with the latter being documented for the first time. Both *M. pilosus* and *M. chinensis* are considered regionally threatened. The potential interspecific competition observed in our findings, particularly between *M. pilosus* and *M. chinensis* during wet season, underscores the importance of protecting macroinvertebrate food sources in local terrestrial ecosystem to prevent intensifying competition among the congenerics. To safeguard the sustainability of vulnerable *M. pilosus*, it is also crucial to preserve the integrity of marine, brackish, and freshwater ecosystems in Hong Kong, as these diverse environments serve as essential foraging grounds for *M. pilosus* to capture the diverse range of fish species.

## 5. Conflict of Interest

The authors declare that the research was conducted in the absence of any commercial or financial relationships that could be construed as a potential conflict of interest.

## 6. Data Availability Statement

Raw sequencing data will be uploaded to the NCBI Sequence Read Archive (SRA). Accession number will be provided after the manuscript is accepted.

## Author contributions

S.Y.W.S. Conceptualization; E.S.K.P. and S.Y.W.S. Methodology; E.S.K.P., J.C.T.C., D.T.C.C., C.T.S., W.C.T., and S.Y.W.S. Investigation; X.W. Formal analysis; C.T.S., W.C.T., and S.Y.W.S. Resources; X.W. Visualization; X.W. and E.S.K.P. Writing - Original Draft; E.S.K.P., S.Y.W.S. and all authors Writing - Review & Editing; S.Y.W.S. Supervision; S.Y.W.S. Project administration; S.Y.W.S. Funding acquisition.

## Statement on ethics

Approvals for animal experiments were granted by the Department of Health (ref. 19- 177 in DH/SHS/8/2/3 Pt. 30), the University of Hong Kong (ref. 4963-19), and the Agriculture, Fisheries, and Conservation Department (AFCD; ref. 35 in AF GR CON 09/51 Pt.8).

## Acknowledgments

This study was funded by the Environment and Conservation Fund (ref. ECF 94/2019) in Hong Kong, China. We express our gratitude to the AFCD of the HKSAR Government for their generous support. Special thanks are extended to Chi Pan Tong and Alex Wai Kit Lo for their assistance during our study. The High-Performance Computing (HPC) service provided by the Information Technology Services at the University of Hong Kong supported our data analysis.

Supplementary Materials

## Supplementary Materials and Methods

### Preparation of DNA metabarcoding libraries through 2-step PCRs

For 18S rDNA libraries, 1^st^ step PCRs were carried out in 30 μl reactions containing 6 μl of 5X GoTaq Flexi Buffer, 0.6 μl of 10 mM dNTP Mix, 3.6 μl of 25 mM MgCl_2_, 0.15 μl of 5 U/μl GoTaq G2 Flexi DNA Polymerase, 1 μl of extracted DNA, 0.6 μl of each assigned 10 μM forward and reverse primer uniquely tagged with heterogeneity spacer (Cruaud et al. 2017), 6 μl of 10 % DMSO (Sigma), 0.15 μl of 20 mg/ml BSA (NEB) and ultrapure water. Thermal cycling condition was 95 °C for 2 min; 25 cycles of 95 °C for 30 sec, 50 °C for 30 sec and 72°C for 30 sec; and final extension at 72 °C for 5 min.

For COI libraries, the 1^st^ step PCRs were carried out in 6 μl of 5X GoTaq Flexi Buffer, 0.6 μl of 10 mM dNTP Mix, 3.6 μl of 25 mM MgCl_2_, 0.15 μl of 5 U/μl GoTaq G2 Flexi DNA Polymerase, 1 μl of extracted DNA, 1.5 μl of each assigned 10 μM forward and reverse primer uniquely tagged with heterogeneity spacer (Cruaud et al. 2017), 6 μl of 10 % DMSO (Sigma), 0.15 μl of 20 mg/ml BSA (NEB) and ultrapure water. Thermal cycling condition was 95 °C for 2 min; 25 cycles of 95 °C for 30 sec, 48 °C for 30 sec and 72 °C for 30 sec; and final extension at 72 °C for 5 min. Target PCR products generated from COI primers were gel extracted to eliminate non-targets.

For 12S rDNA libraries, 1^st^ step PCRs were carried out in 30 μl reactions containing 6 μl of 5X GoTaq Flexi Buffer, 0.6 μl of 10 mM dNTP Mix, 3.6 μl of 25 mM MgCl_2_, 0.15 μl of 5 U/μl GoTaq G2 Flexi DNA Polymerase, 1 μl of extracted DNA, 0.6 μl of each assigned 10 μM forward and reverse primer uniquely tagged with heterogeneity spacer (Cruaud et al. 2017), 6 μl of 10 % DMSO (Sigma), 0.15 μl of 20 mg/ml BSA (NEB) and ultrapure water. Each of the 1^st^ step PCR products was eluted in 20 μl elution buffer and used as the templates for the 2^nd^ step PCRs.

The 2^nd^ step PCRs of both markers were performed with the same conditions: 2^nd^ step PCRs were performed in 45 μl reactions comprising 9 μl of 5X GoTaq Flexi Buffer, 0.9 μl of 10 mM dNTP Mix, 5.4 μl of 25 mM MgCl2, 0.225 μl of 5 U/μl GoTaq G2 Flexi DNA Polymerase, 20 μl of 1^st^ step PCR products, 0.9 μl of each assigned 10 μM forward and reverse index primers, 4.5 μl of 10 % DMSO (Sigma) and ultrapure water. PCR condition was 95 °C for 2 min; 15 cycles of 95 °C for 30 sec, 55 °C for 30 sec, and 72 °C for 30 sec; and final extension at 72 °C for 5 min. We used the PureLink PCR Purification Kit (Invitrogen, Carlsbad, CA) to clean up all PCR products. The band sizes of all ^1st^ and 2^nd^ PCR products were checked on 1.5% agarose gels by gel electrophoresis.

## Supplementary Figures

**Figure S1.**
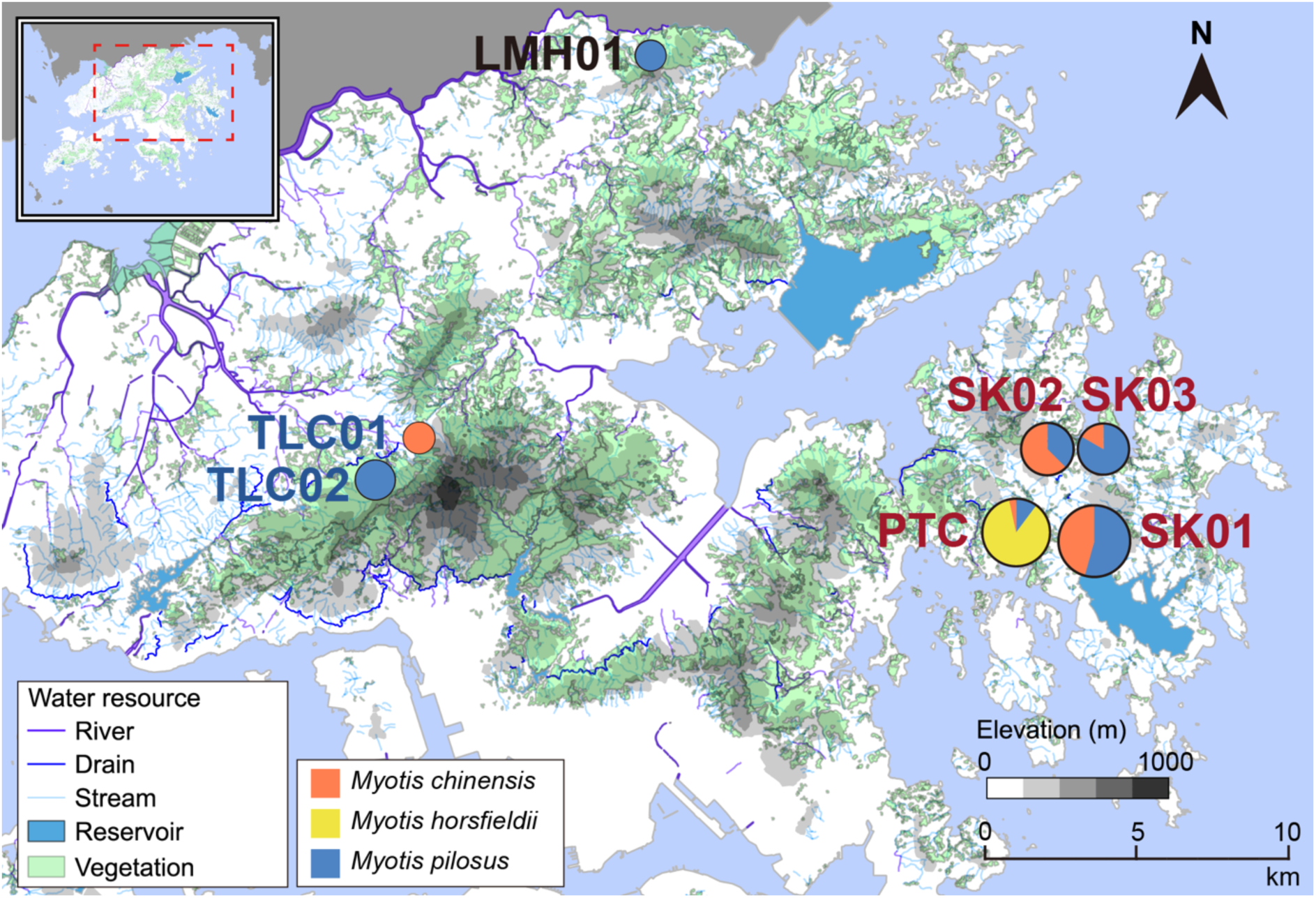
The fecal samples of *Myotis* bats were collected from seven sites in Hong Kong. The small map at the upper left corner represents Hong Kong at the south coast of China, with the area enclosed in a red box magnified for ease of viewing. The pie charts in the magnified map represent the relative quantity (not to scale) of samples collected at each location, with larger pie charts indicating a larger sample size. Sites in close proximity were grouped as sampling locations for analyses (see Table S1). The locations include LMH (indicated in black), TLC (indicated in dark blue, including TLC01 and TLC02), and SK (indicated in dark red, including PTC, SK01, SK02, and SK03). LMH, Lin Ma Hang. TLC, Tai Lam Chung. PTC, Pak Tam Chung Camp. SK, Sai Kung.

**Figure S2.**
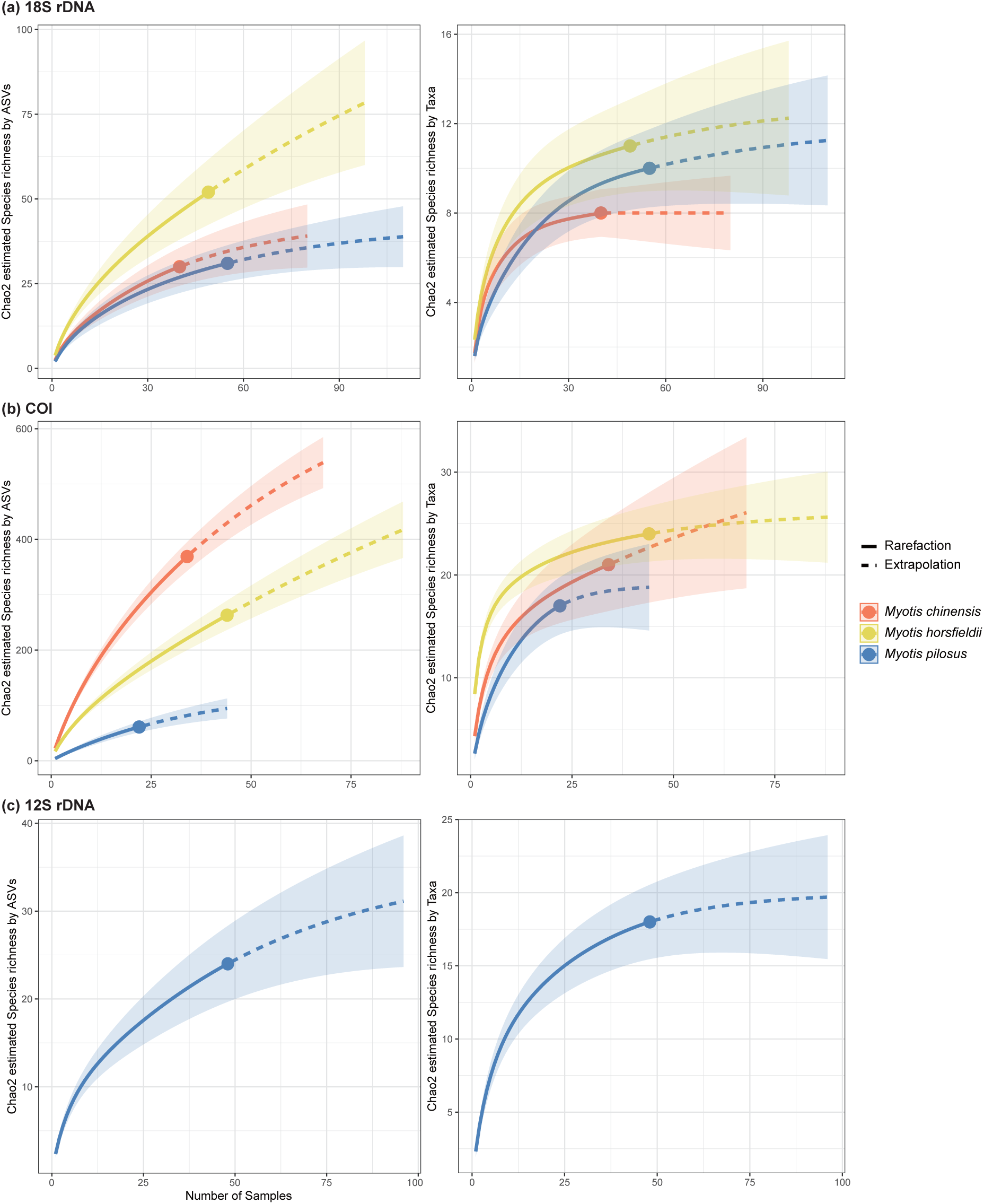
Rarefaction curves based on Chao2 species richness. The Chao2 species richness was estimated from hill numbers of diversity order *q*=0, which were evaluated based on ASVs (left panel) or taxa (b) in each *Myotis* species. Dietary ASVs or taxa were identified by (a) 18S rDNA, (b) COI, and (c) 12S rDNA marker gene. The shading area represent the 95% confidence intervals, which were estimated using a bootstrap method based on 100 replicates.

**Figure S3.**
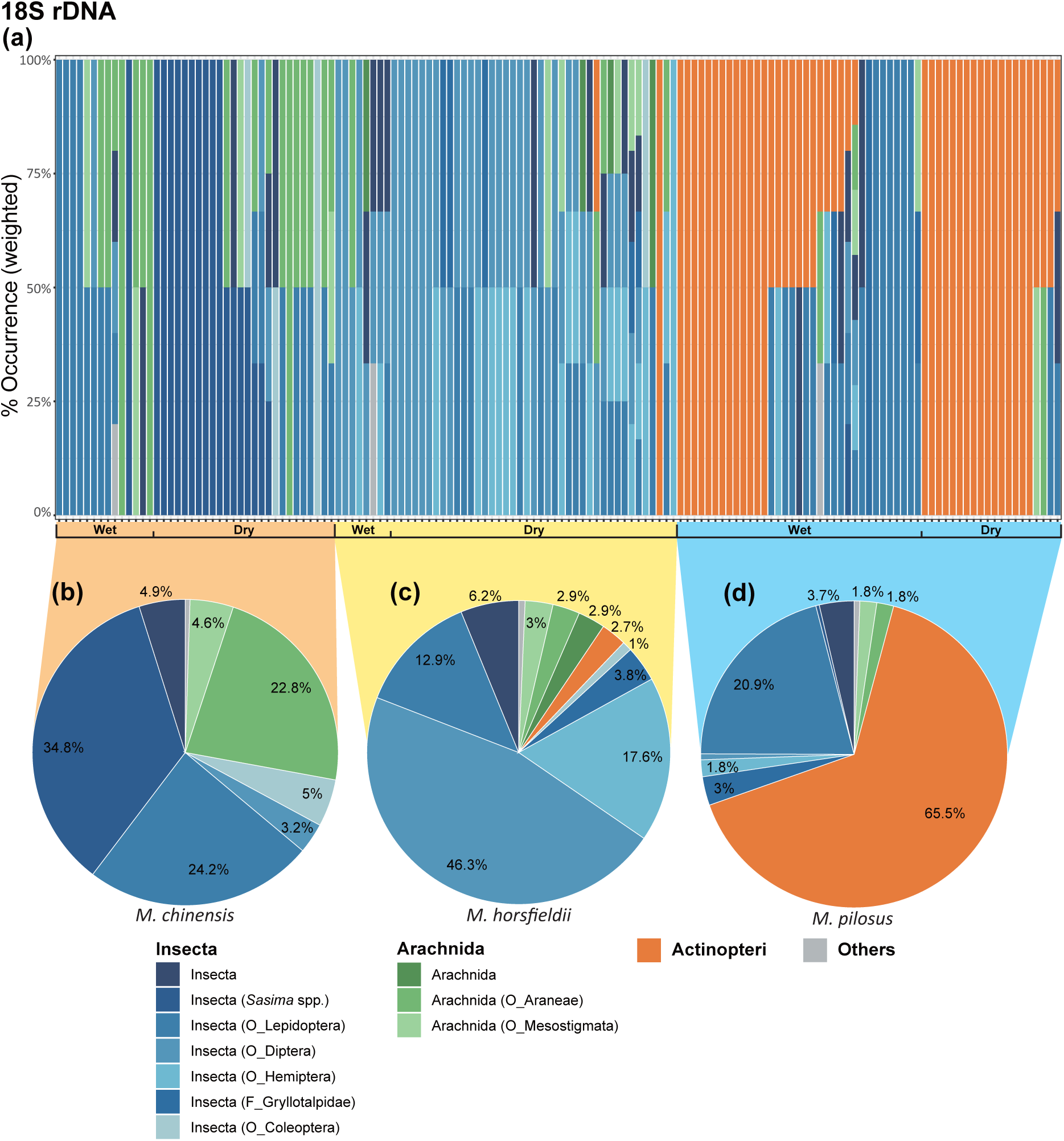
Dietary composition of three *Myotis* bat species based on taxa detected in fecal samples using the 18S rDNA marker. The weighted percentages of occurrence of each taxon were visualized at both (a) bat individual level (n=144) and (b-d) bat species level for (b) *M. chinensis* (n=40), (c) *M. horsfieldii* (n=49), and (d) *M. pilosus* (n=55). Prey in classes with read abundance lower than 0.1% of all taxa detected were grouped into “Others”. Only taxa with abundance higher than 1% in relative read abundance are shown. F, Family. O, Order.

**Figure S4.**
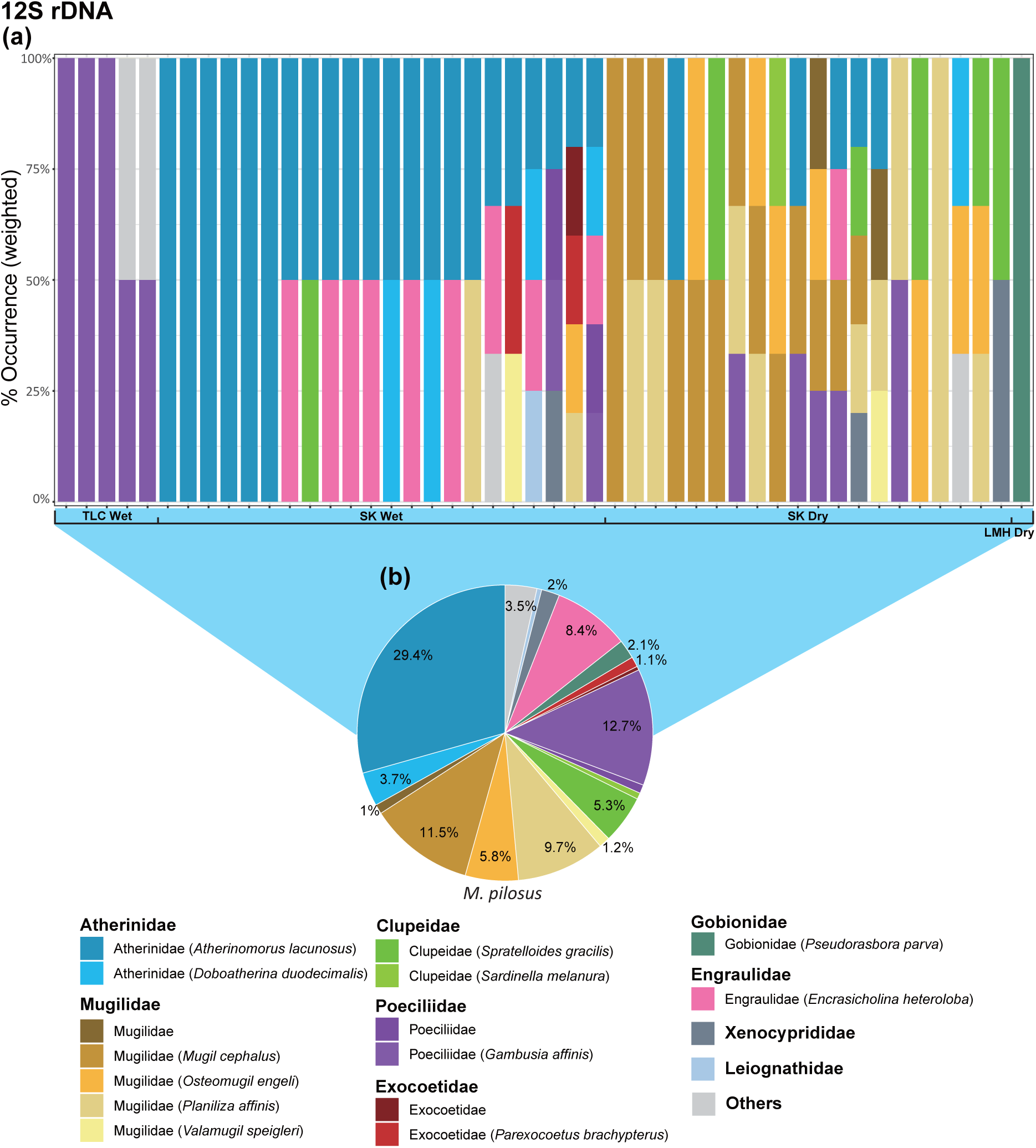
Dietary composition of *Myotis pilosus* based on taxa detected in fecal samples using the 12S rDNA marker. The weighted percentages of occurrence of each taxon were visualized at both (a) individual (n=48) and (b) species levels. Individual bars were grouped by sampling locations and seasons. Prey in classes with read abundance lower than 0.1% of all taxa detected were grouped into “Others”. Only taxa with abundance higher than 1% in relative read abundance are shown. SK, Sai Kung (including Pak Tam Chung); TLC, Tai Lam Chung; LMH, Lin Ma Hang. Refer to Fig. S1 for locations.

**Figure S5.**
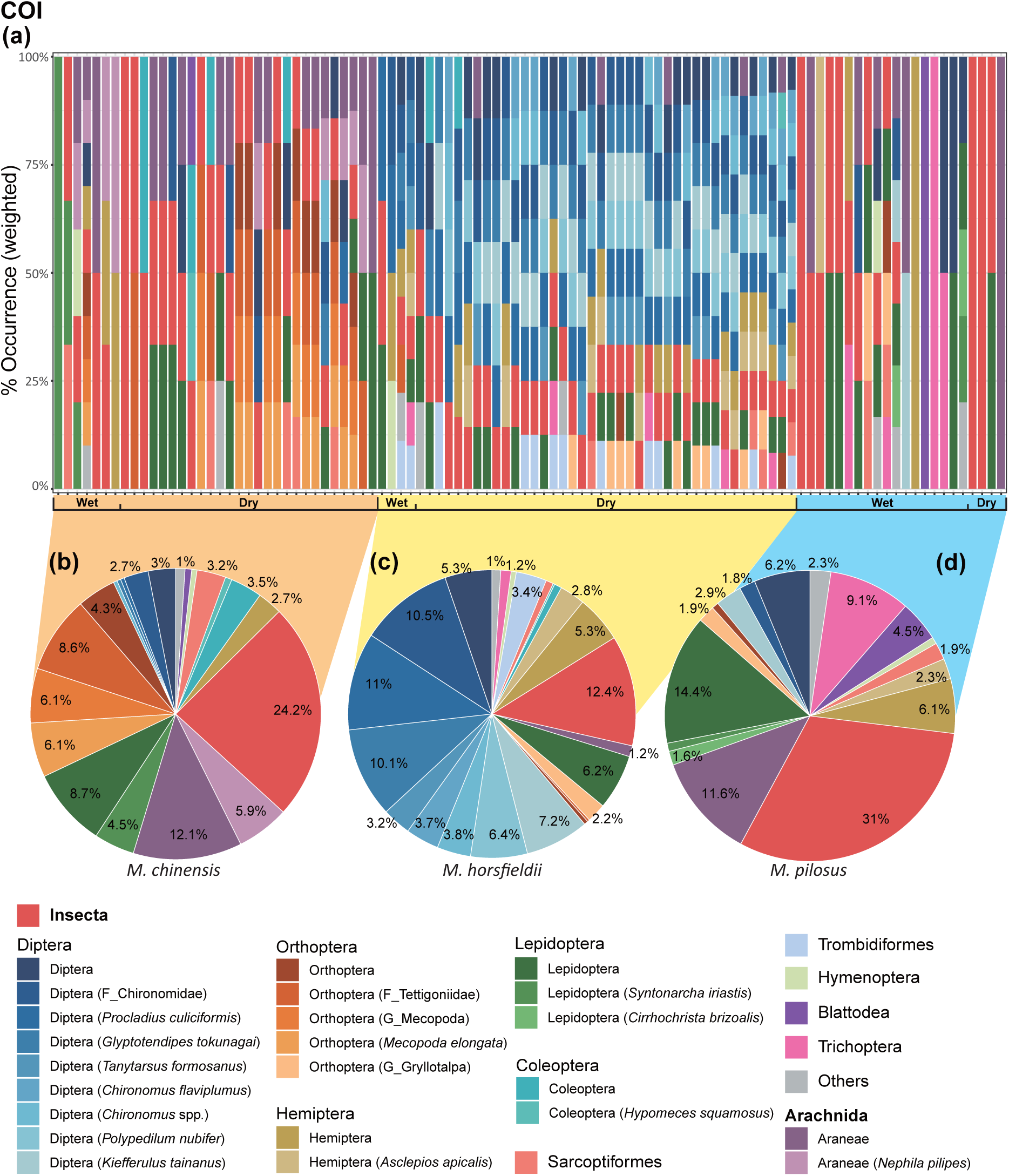
Dietary composition of three *Myotis* bat species based on taxa detected in fecal samples using the COI marker. The weighted percentages of occurrence of each taxon were visualized at both (a) individual level (n=102) and (b-d) species level for (b) *M. chinensis* (n=34), (c) *M. horsfieldii* (n=44), and (d) *M. pilosus* (n=22). Prey in classes with read abundance lower than 0.1% of all taxa detected were grouped into “Others”. Only taxa with abundance higher than 1% in relative read abundance are shown. G, Genus. F, Family.

**Figure S6.**
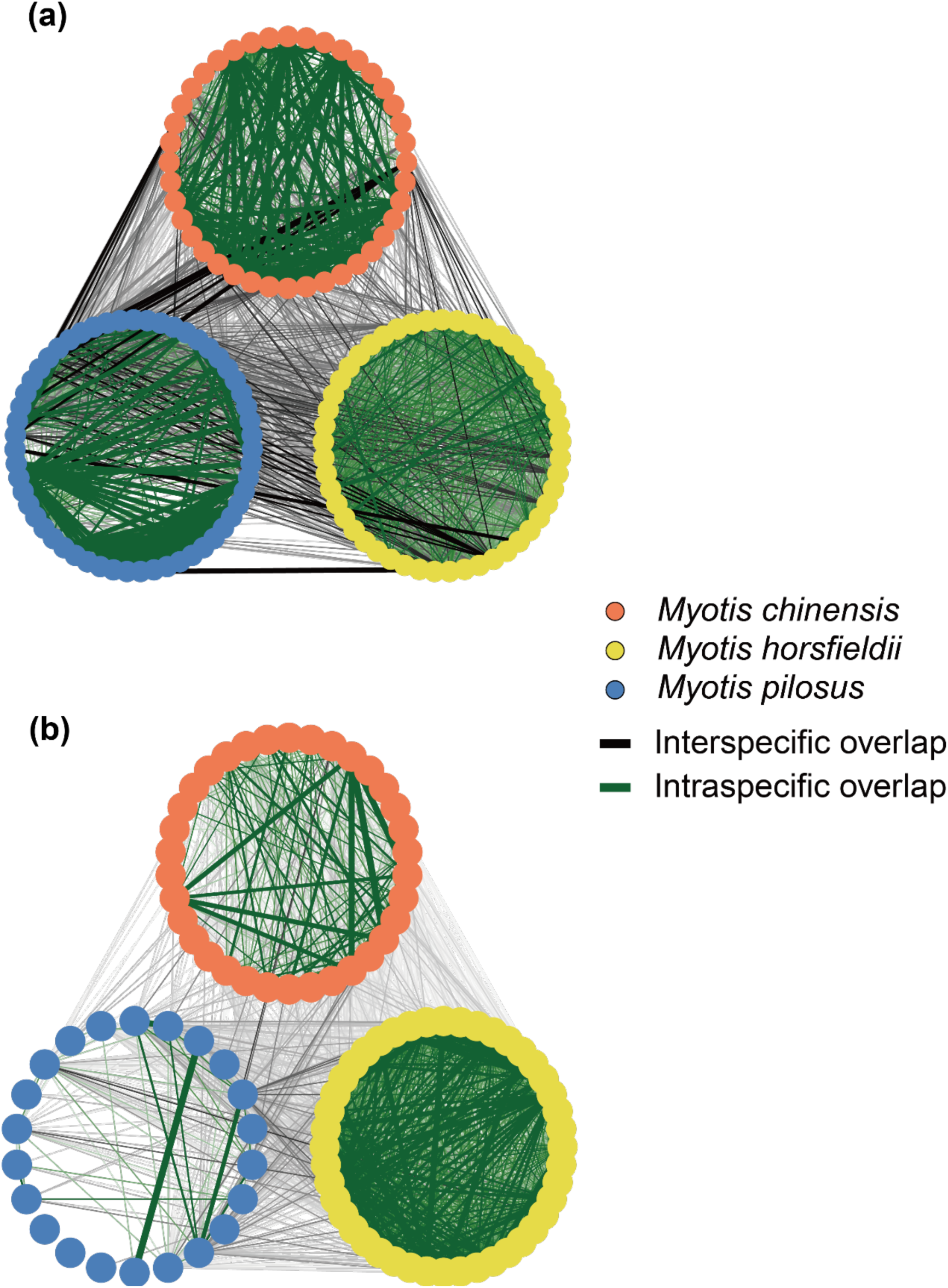
Dietary overlap of three sympatric *Myotis* bat species. Each colored circle represents a single bat sample. Dietary overlap between individual pairs was calculated based on their diet composition determined by (a) 18S [*M. chinensis* (n=40), *M. horsfieldii* (n=49), and *M. pilosus* (n=55)] and (b) COI [*M. chinensis* (n=34), *M. horsfieldii* (n=44), and *M. pilosus* (n=22)] markers. The thickness of the line connecting circles represents the extent of dietary overlap between individuals.

## Notes

### Competing Interest Statement

The authors have declared no competing interest.

### Summary of Updates

New analysis added; Figures revised; and supplementary files updated.

